# Spatial distribution of blood-brain barrier membrane proteins is controlled by sorting motifs and physiological signals in vivo

**DOI:** 10.64898/2026.05.10.724157

**Authors:** Joseph Amick, Shamika Bhandarkar, Douglas R. Wilcox, Solenn Boloker, Yi Daniel Lu, Chenghua Gu

## Abstract

The blood-brain barrier (BBB), formed by brain endothelial cells (BECs), creates a safe and homeostatic environment for proper brain function. Together with pericytes and astrocytes, the BBB controls substance influx and efflux into and out of the brain. While BECs are extraordinarily thin, their luminal and abluminal plasma membranes face markedly different environments: the blood and brain parenchyma, respectively. How BBB membrane proteins are spatially distributed between these membranes and the factors regulating their localization are not clear. Here, we establish a method for measuring polarized protein sorting at the BBB *in vivo*. We characterize the distribution of transporters and receptors on the luminal and abluminal plasma membranes and identify protein-intrinsic motifs that control polarized sorting. Finally, we observe a change in membrane protein distribution that aligns with circadian rhythms, revealing an under-appreciated dimension of BBB dynamics. This method can be broadly used for evaluating membrane protein landscape changes during development and disease and for choosing optimal molecular targets for BBB-crossing therapeutics.

## Introduction

The blood-brain barrier (BBB) is the interface that tightly controls molecular trafficking between the blood and brain to maintain a homeostatic environment for neuronal health and function^1^. Endothelial cells, which form the inner lining of blood vessels, are the anatomical site of the BBB^2^. In addition to controlling paracellular and transcellular trafficking, brain endothelial cells (BECs) express a multitude of transporters for nutrient delivery and waste clearance. An intact BBB is a major obstacle for central nervous system (CNS) drug delivery^3^, whereas impaired BBB function is observed in neurodevelopmental and neurodegenerative disease^4–8^. Thus, therapeutic strategies to manipulate the barrier for different purposes are needed. Such strategies hinge on greater understanding of signaling and transport pathways in BECs, which are governed by membrane proteins.

BECs are polarized^9^. They possess a distinct luminal plasma membrane that faces the blood, in contact with humoral factors, and an abluminal plasma membrane that faces the brain parenchyma and interacts with brain cells and brain-derived factors^10^. A remarkable challenge for BECs is that their luminal and abluminal plasma membranes must interact with these different environments while being separated by a tiny distance; most of the BEC is just 200 nm thick^11^. BECs interact with these different environments in part via their membrane proteins. Thus, targeting membrane proteins to the right locations is likely to be critical for BBB integrity, nutrient delivery, and removal of toxic substances. Although transcriptomic studies of BECs have exponentially increased in recent years, yielding a list of genes uniquely expressed at high levels in BECs^12–14^, much less is known about how these proteins are distributed spatially. Still less is known about the rules and molecular machinery that govern polarized membrane protein sorting in BECs.

Several obstacles have limited progress in understanding BEC membrane protein spatial distribution. First, the thin dimensions of BECs challenge the diffraction limit of light microscopy. Previous approaches have primarily relied on immuno-electron microscopy, but very few antibodies are compatible with this method and the nature of labeling is extremely sparse, limiting its quantitative potential. Second, neighboring cells like pericytes and astrocytes tightly associate with BECs within the complex brain environment, hindering the ability to distinguish between BEC, pericyte or astrocyte membranes^15–18^. Third, *in vivo* analysis of BECs is critical; *ex vivo*, BECs rapidly de-differentiate, losing expression of key BBB proteins after purification from the brain^19^.

To overcome these limitations, we developed a method leveraging the unique anatomy of BECs, super-resolution microscopy, and new genetic tools to quantitatively reveal luminal and abluminal protein distribution in BECs *in vivo*. We applied this approach to measure the relative levels of a variety of transporters and receptors on the luminal and abluminal plasma membranes, identifying subsets of these proteins enriched at each surface to different degrees. Moreover, we identified protein-intrinsic motifs that control BBB membrane protein polarized sorting. Finally, we examined membrane protein spatial distribution between morning and night and identified a shift in CD31 localization, revealing a surprising new facet of BEC dynamics. This method has broad applications, including further understanding of fundamental BBB cell biology, evaluation of changes of distribution of membrane proteins during development and disease states, and insight into optimal molecular targets for BBB therapeutics.

## Results

### Development of an approach to image and measure luminal-abluminal membrane protein distribution in BECs *in vivo*

We set out to measure how proteins are distributed between the luminal and abluminal plasma membranes *in vivo*. The small distance separating the luminal and abluminal plasma membranes renders them difficult to resolve by light microscopy. However, as is apparent under electron microscopy, in the area surrounding the endothelial cell nucleus (Fig. 1A) the two plasma membranes are separated by a distance that can be resolved by stimulated emission depletion super-resolution microscopy (STED). Taking advantage of this anatomical feature, we used STED to image this area of the endothelial cell in mouse brain sections immunostained by different membrane protein antibodies. To quantitively measure polarized protein distribution, the immunofluorescent signal at luminal and abluminal plasma membranes was measured in the area surrounding the endothelial nucleus. Then, the abluminal enrichment for a given protein was calculated as the mean abluminal intensity divided by the mean luminal intensity (>1 indicates abluminal enrichment, 1 indicates equal distribution, <1 indicates luminal enrichment). The percentage of the total protein which was luminal or abluminal was also calculated (Fig. 1B).

**Figure 1.**
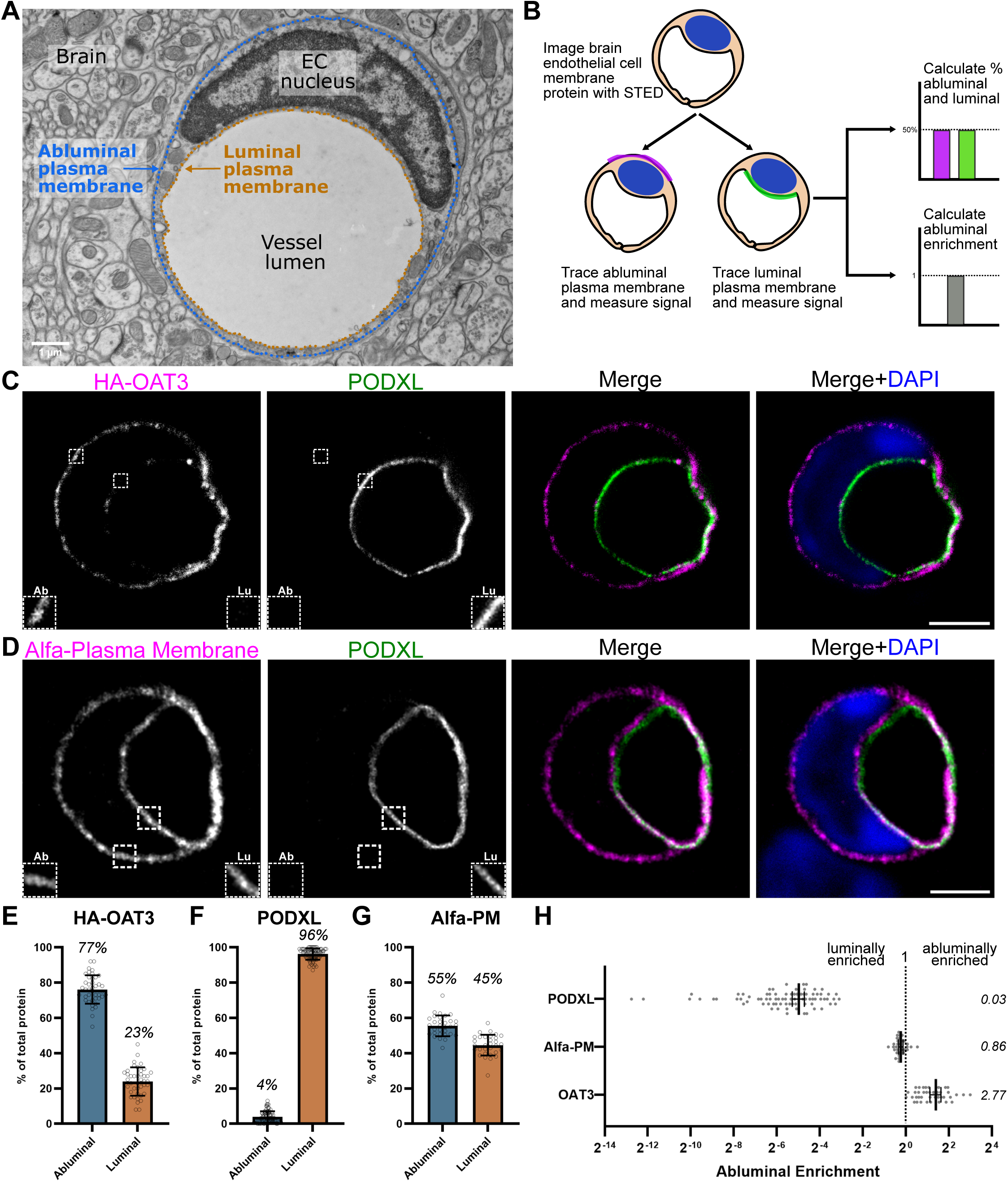
A super-resolution strategy for measuring polarized protein sorting in brain endothelial cells in vivo. **(A)** Electron micrograph of a mouse brain endothelial cell (BEC). The positions of the luminal plasma membrane (orange) and abluminal plasma membrane (blue) are indicated. **(B)** Schematic diagram illustrating the approach for measuring and calculating the percentage distribution at the luminal and abluminal plasma membrane and its abluminal enrichment. **(C)** STED image of AAV-BI30-transduced BEC expressing HA-tagged OAT3 co-stained with luminal plasma membrane marker PODXL. Left inset: abluminal membrane (Ab). Right inset: luminal membrane (Lu). Nucleus is labeled by DAPI (4′,6-diamidino-2-phenylindole). **(D)** STED image of AAV-BI30-transduced BEC expressing an Alfa-tagged general plasma membrane marker (Alfa-PM) co-stained with PODXL. **(E-G)** Quantification of the percentage of HA-OAT3, PODXL and Alfa-PM at the luminal and abluminal plasma membranes (mean ± SD). Datapoints are measurements from individual cells. Mean luminal and abluminal percentages are listed. **(H)** HA-OAT3, PODXL and Alfa-PM abluminal enrichments (mean ± 95% confidence interval). Mean abluminal enrichments are listed. Scale bars: 2µm.

We began using this strategy to measure known luminal and abluminal proteins, podocalyxin (PODXL) and OAT3, respectively. PODXL is a cell surface sialomucin expressed at the apical membrane of podocytes^20^. It is also expressed in endothelium in a variety of organs including brain and was localized to the luminal membrane by immunoelectron microscopy^21^. Indeed, our STED imaging of PODXL distribution revealed it was 96.1% luminal with an abluminal enrichment of 0.03 (Fig. 1C,F,H). OAT3, an organic anion transporter, was reported to localize to the endothelial cell abluminal membrane in rat brains^22,23^. However, at the transcript level, in addition to being highly expressed in BECs, OAT3 expression has been detected in brain pericytes, which could confound the analysis^14^. Additionally, we could not identify specific antibodies against OAT3 for immunohistochemistry in mice. To selectively visualize and measure OAT3 in BECs, we used the BI30 capsid, a systemically-delivered AAV that efficiently transduces endothelial cells in the mouse CNS^24^, to express an HA-tagged OAT3 in mouse BECs *in vivo* (Fig. 1C). In contrast to PODXL, HA-OAT3 was 76.9% abluminal with an average abluminal enrichment of 2.77 (Fig. 1E,H). Finally, we designed a construct to non-selectively label both the luminal and abluminal plasma membranes by fusing the CAAX-box membrane targeting motif from K-Ras to the Alfa epitope tag for detection, which we expected to have a roughly equal distribution between the two membranes^25,26^. Indeed, this construct was 55.5% abluminal and had an abluminal enrichment of 0.86 (Fig 1D,G,H). (The longer trace along the abluminal membrane than the luminal membrane pushes the abluminal percentage slightly higher than 50% when intensities on each side are similar.) These results served as a proof of concept that this approach could reliably measure the distribution of membrane proteins in BECs *in vivo*.

### Spatial distribution of BEC membrane proteins in mouse and human

We next used this approach to measure the spatial distribution of a panel of BEC membrane proteins, including both BBB-enriched transporters and receptors, as well as pan-endothelial membrane proteins.

Some proteins were predominantly localized to the luminal membrane. P-glycoportein (P-gp) (98.2% luminal, abluminal enrichment 0.02), a highly expressed protein in BECs, is a broad-specificity ATP-driven pump which extrudes xenobiotics and endogenous hormones from cells, limiting their entry into the brain (Fig. 2A)^27^. Intercellular Adhesion Molecule 2 (ICAM2) (86.5% luminal, abluminal enrichment 0.11) is a constitutively-expressed cell adhesion molecule in endothelial cells that interacts with leukocyte integrins (Fig. 2B)^28^. Basigin (79.1% luminal, abluminal enrichment 0.21) is a glycosylated type I transmembrane protein in the immunoglobulin superfamily highly expressed in BBB-forming brain endothelial cells (Fig. 2C)^12,29^. CD31 (68.9% luminal, abluminal enrichment 0.39) is a pan-endothelial membrane protein that regulates leukocyte transendothelial migration and other processes (Fig. 2D)^30^.

**Figure 2.**
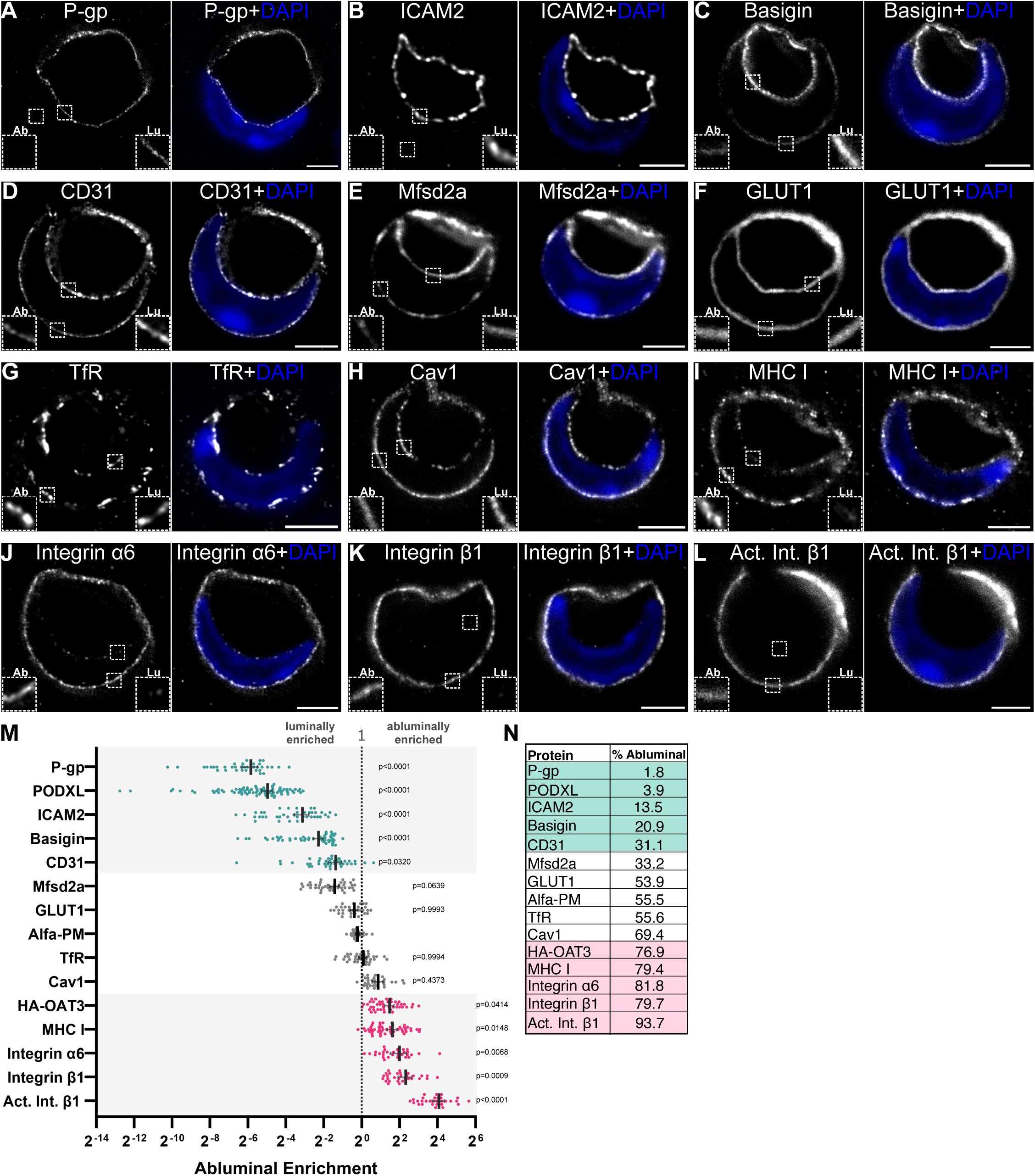
Membrane protein distribution in brain endothelial cells in vivo. **(A-L)** STED images of the indicated proteins with and without DAPI to show the position of the nucleus. Scale bars: 2µm. **(M)** Abluminal enrichment for proteins in A-L (mean ± 95% confidence interval). Datapoints are measurements from individual cells. Proteins were categorized as luminally enriched (teal) or abluminally enriched (pink) if their log-transformed abluminal enrichments differed significantly from the non-targeted control, Alfa-PM, by nested ANOVA with Dunnett’s multiple comparisons test; p-values indicated. **(N)** Summary table of the mean abluminal percentages of the proteins in A-L.

Some proteins showed a more equal distribution between the luminal and abluminal plasma membranes. Mfsd2a (66.8% luminal, abluminal enrichment 0.37) is a transporter of unsaturated phospholipids required for the transport of omega-3 fatty acids into the brain and maintains a unique lipid composition of BECs that suppresses caveolae-mediated transcytosis (Fig. 2E)^31,32^. The glucose transporter GLUT1 (53.9% abluminal, abluminal enrichment 0.76) is highly expressed in BECs and is responsible for transport of glucose into the brain (Fig. 2F)^33^. The transferrin receptor (TfR) (55.6% abluminal, abluminal enrichment 1.05) is also highly expressed in BECs, is critical for the transport of iron into the brain^34^, and is a major target for therapeutic delivery into the brain (Fig. 2G)^35^. Caveolin-1 (Cav1) (69.4% abluminal, abluminal enrichment 1.82) is a membrane scaffolding protein required for caveolae vesicle formation and regulates T cell trafficking across the BBB (Fig. 2H)^36–39^.

Certain proteins displayed abluminal enrichment. BECs express major histocompatibility complex class I (MHC I) proteins, a key group of immune system receptors^14,40^. MHC I abluminal enrichment was 3.07 (79.5% abluminal) (Fig.2I). Integrins are heterodimeric transmembrane proteins that mediate cell-cell and cell-extracellular matrix adhesion; deletion of certain EC-expressed integrins causes BBB defects^41,42^. Integrin α6 (81.8% abluminal, abluminal enrichment 3.97) is a receptor for laminins (Fig. 2J)^43,44^. Integrin β1 showed an abluminal enrichment of 4.97 (79.7% abluminal) (Fig. 2K). Interestingly, an antibody that recognizes the active conformation of integrin β1 showed a further enrichment at the abluminal plasma membrane (16.76 abluminal enrichment, 93.7% abluminal) (Fig. 2L).

For some abluminally-enriched proteins, a potential complication of imaging brain tissue is that BECs form close contacts with pericytes and astrocyte endfeet. Of the proteins we examined, integrin α6 has been detected at the transcript level in astrocytes (at a lower level than BECs) and integrin β1 is expressed in pericytes at high levels^14^. As this had the potential to confound our measurements, we modified our approach or performed additional imaging for these proteins (see methods; Supplemental Figure 1).

Due to the BBB’s therapeutic and disease relevance, we tested if our approach could be applied to human brain tissue samples. Using postmortem brain tissue, we found GLUT1 showed a roughly equal distribution between the luminal and abluminal membranes in human BECs (Supplemental Fig.2A,D). Cav1 was also readily detected on both membranes, though modestly more abluminally-enriched than GLUT1 (Supplemental Fig.2B,D). In contrast, PODXL exhibited a more luminally-enriched distribution than GLUT1 and Cav1 (Supplemental Fig.2C,D). Therefore, human BECs exhibit polarity and different membrane protein distributions can be measured in human brain samples through this approach.

Overall, we observed a broad spectrum of protein distributions (Fig. 2M,N and Supplemental Table 4). To help categorize these different distributions, we statistically compared the abluminal enrichments of the proteins measured above to the control non-targeted plasma membrane marker, Alfa-PM. Proteins whose abluminal enrichments differed significantly from Alfa-PM were categorized as luminally enriched (P-gp, PODXL, ICAM2, basigin and CD31) or abluminally enriched (HA-OAT3, MHC I, integrin α6, integrin β1 and active integrin β1). Proteins whose abluminal enrichments did not differ significantly from Alfa-PM were categorized as non-polarized (Mfsd2a, GLUT1, TfR and Cav1) (Fig. 2M). Importantly, this taxonomy should be viewed as a continuum rather than an absolute classification, reflecting differences that are detectable from the non-polarized control at the sample sizes examined. For example, Mfsd2a and CD31 exhibit similar abluminal enrichments and have p-values of 0.0639 and 0.0320, respectively, placing them just on either side of the significance threshold. In sum, these results indicate the existence of powerful sorting programs in BECs that can produce robust luminal or abluminal enrichment and suggest that diverse sorting mechanisms may operate in BECs.

### Protein-intrinsic motifs control BBB membrane protein localization

We next asked what factors determine the polarized protein distribution in BECs. Much of what is known about mechanisms of polarized membrane protein sorting has been learned from studies of apical-basolateral sorting in cultured epithelial cells and somatodendritic-axonal sorting in cultured neurons^45,46^. While apical determinants tend to be nebulous, some basolateral and somatodendritic determinants are distinct, specific amino acid sequences in the cytoplasmic domains of transmembrane proteins. Such sorting signals include YXXØ and [DE]XXXL[LI], where amino acids are represented by their single-letter code, X represents any amino acid and Ø represents a bulky hydrophobic amino acid. These motifs are recognized by adaptor protein complexes, heterotetramers that interact with both these sequences and clathrin, which couples cargo selection and vesicle formation, promoting basolateral/somatodendritic sorting of motif-containing cargo proteins^47^. Whether these motifs regulate polarized protein distributions in BECs is currently unknown. To identify candidate sorting motifs in BEC-expressed proteins, we computationally searched 15,813 BEC-expressed genes, filtered them based on the presence of a transmembrane domain and their topology to identify YXXØ and [DE]XXXL[LI] sequences in their cytoplasmic regions (Supplemental Table 3). This bioinformatic approach yield a list of membrane proteins containing potential sorting motifs which could promote abluminal enrichment. One such motif, YPTV, was found in OAT3 (Fig. 3A). To test if this sequence indeed is responsible for targeting OAT3 to the abluminal membrane, we generated a Y437A/V440E mutant and expressed the wild type and mutant forms of HA-tagged OAT3 *in vivo* in BECs using the BI30 AAV (Fig. 3B). Remarkably, HA-OAT3’s abluminal enrichment was lost in the Y437A/V440E mutant, which was roughly equally distributed between the luminal and abluminal plasma membranes (Fig. 3B,C,D).

**Figure 3.**
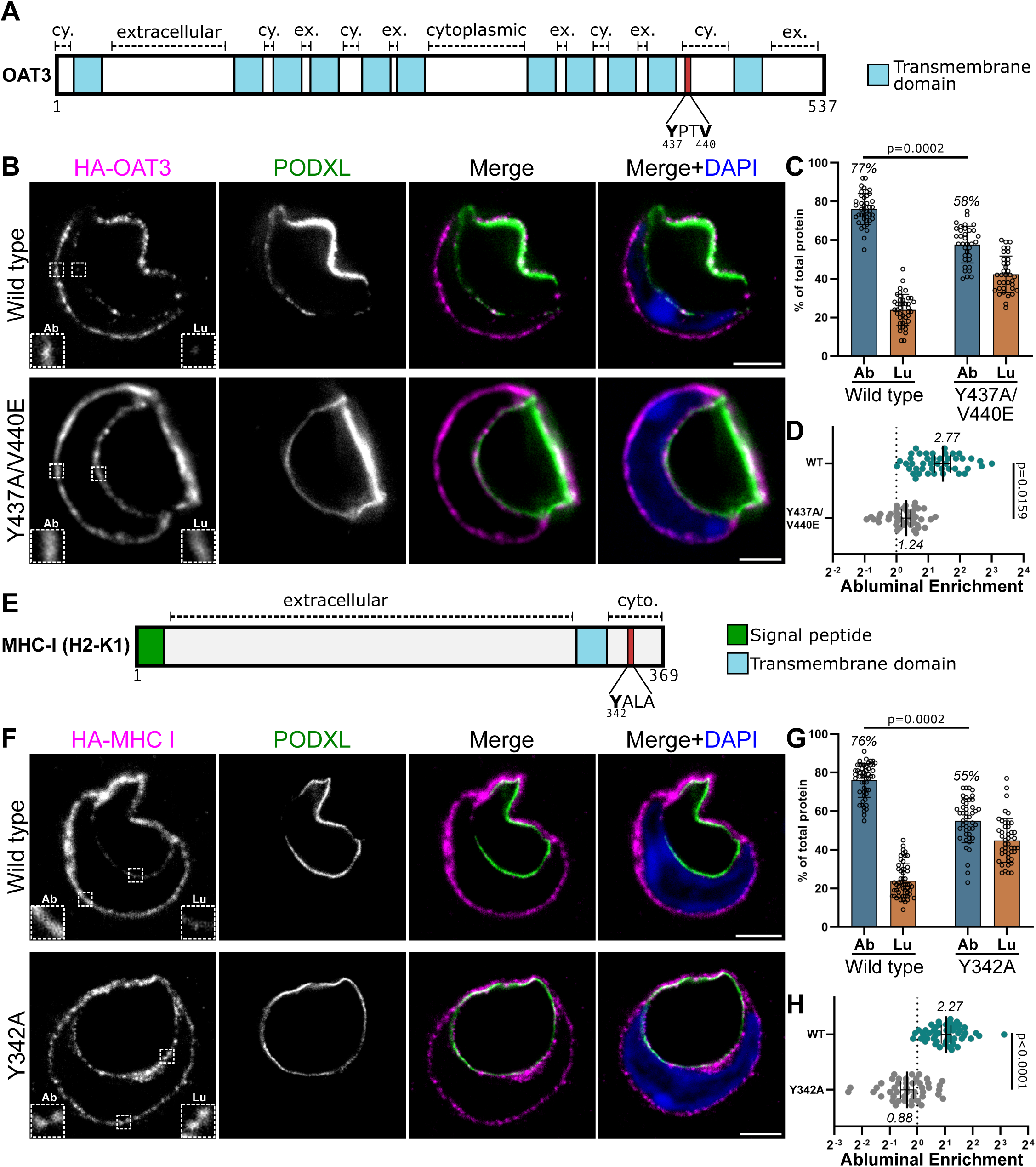
Tyrosine-based sorting motifs promote abluminal plasma membrane enrichment. **(A)** Schematic diagram illustrating OAT3’s topology and the position of tyrosine 437 and valine 440. **(B)** STED images of AAV-BI30 transduced BECs expressing HA-tagged wild type or Y437A/V440E mutant OAT3. Scale bars: 2µm. **(C)** Abluminal (Ab) and luminal (Lu) percentages of wild type and mutant OAT3. p-Value from nested t-test indicated. **(D)** Abluminal enrichment of wild type and mutant OAT3. Nested t-test. **(E)** Schematic diagram of MHC I illustrating the position of tyrosine 342. **(F)** STED images of AAV BI30-transduced BECs expressing HA-tagged wild type or Y342A mutant MHC I. Scale bars: 2µm. **(G)** Abluminal and luminal percentages of wild type and mutant MHC I; nested t-test. **(H)** Abluminal enrichment of wild type and Y432A mutant MHC I; nested t-test.

Another abluminally-enriched protein, MHC I, contains a conserved tyrosine residue in its cytoplasmic domain (Fig.3E)^48^. While this YALA sequence is not a canonical YXXØ motif, it was shown to mediate an interaction with adaptor protein complex-1 in dendritic cells^49^. We tested the role of this tyrosine residue in MHC I’s abluminal enrichment by expressing HA-tagged wild type or Y342A mutant MHC I via the BI30 AAV (Fig. 3F). The abluminal enrichment of HA-MHC I was lost in the Y342A mutant HA-MHC I, which showed a roughly equal distribution between the two plasma membranes (Fig. 3F,G,H). Therefore, tyrosine-based motifs are key sorting signals promoting abluminal enrichment in BECs.

Because overexpression of tagged proteins can sometimes influence their localization, we assessed whether HA-tagged wild type MHC I expressed via BI30 localized similarly to endogenous MHC I. Comparing their abluminal enrichments revealed no significant difference between the two groups (Supplemental Fig. 3A), supporting the conclusion that HA-tagged wild-type MHC I exhibits localization comparable to that of the endogenous protein.

Mutations in membrane proteins can result in their misfolding and retention in the endoplasmic reticulum (ER). To examine if the mutant localization reflected ER retention, we used a well-established ER targeting sequence from cytochrome p450^50^ fused to the Alfa epitope tag and expressed this construct via the BI30 virus, allowing us to label the ER in BECs *in vivo* (Supplemental Fig. 3B). The ER in BECs showed a strong enrichment surrounding the nucleus, unlike wild type and mutant OAT3 and MHC I (Supplemental Fig. 3C). Thus, the changes in OAT3 and MHC I mutant localization do not reflect ER retention, indicating that protein-intrinsic sorting motifs regulate abluminal localization in BECs *in vivo*.

### Extrinsic factors modulate membrane protein spatial distribution

Given the emerging recognition of the dynamic properties of BECs in BBB permeability and transport functions, we asked if protein distributions may be dynamically regulated across physiological states, particularly in light of well-established time-of-day differences in cardiovascular function. For example, blood pressure exhibits circadian rhythmicity and the onset of acute cardiovascular events like myocardial infarction, sudden cardiac death or stroke occur most frequently during morning hours^51–54^. At the BBB, circadian changes to efflux transport have been observed^55–58^. Increased permeability of the blood retina barrier has been observed in the evening compared to the morning^59^.

To test this hypothesis, we measured membrane protein distributions in mouse brains harvested during the morning (mouse rest period, zeitgeber time (ZT) 2) and at night (mouse active period, ZT14). CD31, strikingly, shifted significantly from a nearly equal distribution between the luminal and abluminal plasma membranes in the morning to a luminally-enriched distribution at night (Fig. 4A,B). In contrast, GLUT1 (Fig. 4C,D), ICAM2 (Fig. 4E,F), PODXL (Fig. 4G,H), and Cav1 (Fig. 4I,J) did not change significantly. These results indicate that transmembrane protein spatial distribution between luminal and abluminal surfaces can be dynamically regulated, which may contribute to changes in BBB permeability and transport functions observed in various physiological and disease states. Moreover, understanding BBB membrane proteins’ relative spatial distribution, and the conditions that change this distribution, will be critical in the development of therapeutics that target luminal membrane proteins in order to cross the BBB, Thus, this *in vivo* spatial distribution method can be harnessed for a wide variety of applications.

**Figure 4.**
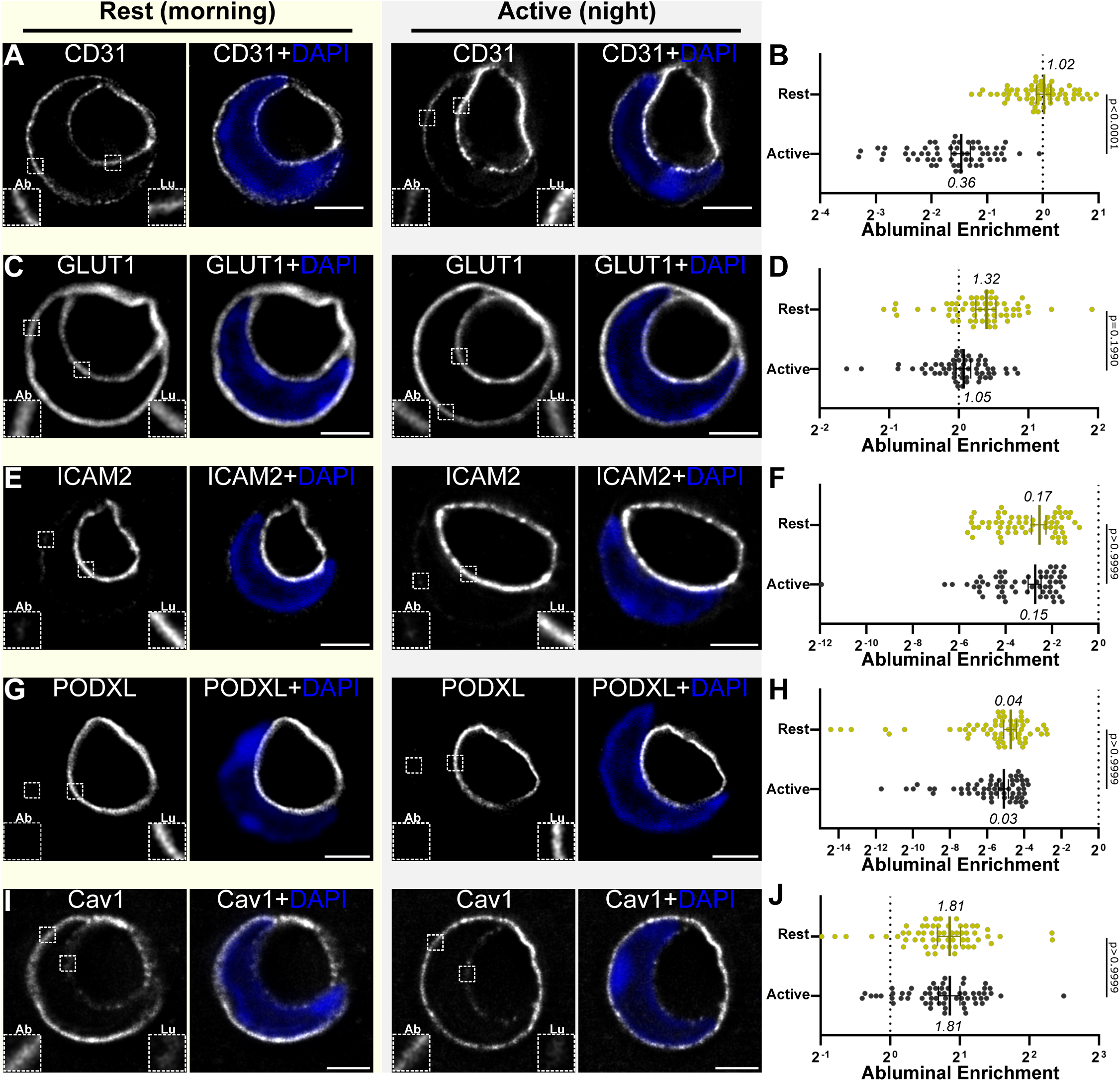
Membrane protein distribution in day versus night. (A,C,E,G,I) STED images of the indicated proteins from mouse brains harvested during the mouse rest (morning, ZT2) or active period (night, ZT14). Scale bars: 2µm. **(B,D,F,H,J)** Graphs of abluminal enrichment during rest and active periods for the indicated proteins (mean ± 95% confidence interval). Mean abluminal enrichments are listed. Nested one-way ANOVA with Bonferroni’s multiple comparisons test; p-values indicated.

## Discussion

In this study, we present an approach to measure the spatial distribution of membrane proteins at the BBB *in vivo*. Because membrane proteins govern fundamental aspects of BEC biology and represent important therapeutic targets, determining their localization to the luminal or abluminal membranes of BECs advances understanding of BBB transport and signaling and informs the viability of candidate receptors for drug delivery. Such measurements are critical to perform *in vivo* because BECs undergo broad changes in their transcriptional profile and lose BBB properties when isolated from brain and cultured *in vitro*^19^; meanwhile, commonly-used BBB models lack key endothelial features^60^. Bringing together super-resolution imaging and recently-developed genetic tools, we uncovered intrinsic motifs within transmembrane proteins that determine their localization in BECs and demonstrate that BBB protein spatial distribution can also be dynamically regulated by extrinsic factors under different physiological conditions. These results open new lines of inquiry into the interactions of the BBB with its environment that were previously not possible.

We identified motifs in OAT3 and MHC I that promote abluminal targeting, shedding light onto one sorting mechanism BECs use to generate different membrane protein distributions. Luminally-enriched proteins may rely on diverse determinants; in epithelial cells, apical sorting determinants are more ambiguous than the discrete amino acid sequences used for basolateral sorting. Biophysical features of apical proteins, including cytoplasmic domain size and a preference for a more tightly packed apical lipid environment, have been implicated in their sorting^61,62^. Understanding luminal protein sorting mechanisms in BECs is a major future direction, as the luminal membrane is the primary site of communication between the BBB and the blood and is thus a major interface of the brain-body axis. Ultimately, this line of work may allow for prediction of protein spatial localization at the BBB, guiding the development of tools that aim to modify or manipulate the BBB.

The brain microenvironment regulates the properties of BECs, including gene expression and BBB permeability. Our approach could also be used to investigate the role of this milieu, including the influence of other cells in the neurogliovascular unit, on BEC protein localization. Collagen type IV alpha 1 chain and PODXL were found to be normally localized in a pericyte-deficient mouse model, indicating there was not a general loss of polarity in the absence of pericytes^63^. However, pericytes or other brain cells may provide cues that influence the spatial distribution of specific transporters and receptors, such as through the regulation of sorting pathways.

Understanding membrane protein distribution in BECs has implications for drug delivery to the brain. BEC membrane proteins have been used in a BBB-crossing strategy to shuttle a therapeutic along with endogenous molecules transported from the blood into the brain. TfR is a major target in this approach, wherein a TfR-binding antibody is coupled to a therapeutic to be delivered to the brain^35,64,65^. Basigin has also been used as a target for brain delivery in a similar way (Zuchero et al., 2016). Efforts to identify BBB receptors that enhance brain delivery of therapeutics are ongoing. The localization of candidate receptors may prove to be a useful characteristic, among many other factors, for evaluating potential targets. For example, a high level of luminal enrichment would make a receptor more abundant in a location where it is accessible for antibody binding after intravenous administration but could indicate little transcytosis. The ability to obtain data from human samples (Supplemental Fig. 2) is particularly relevant for such approaches. This method may also be useful for examining potential changes in brain endothelial polarity in the context of neurological disease and for understanding disease-relevant BBB transport processes. Vascular dementias represent the second most common type of dementia^66^, while epilepsy, multiple sclerosis, glioblastoma, and Alzheimer’s disease, among others, critically involve aspects of neurovascular dysfunction^67^. In Alzheimer’s disease, P-gp and other endothelial transporters have been implicated in Aβ clearance, and in epilepsy altered glucose metabolism and GLUT1 levels have been observed^68–72^. Integrating such observations with information about transporter and receptor polarity could therefore advance understanding of neurovascular dysfunction in disease.

Our unexpected observation of a variable distribution of CD31 between day and night demonstrates protein localization at the BBB is not purely static and a function of protein-intrinsic factors; instead, the distribution between the luminal and abluminal membranes can shift downstream of physiological changes. This change in CD31’s localization opens new questions and opportunities for future investigation. Regarding CD31 specifically, these include the molecular factors that regulate its shift and the functional implications for vascular adaptations between day and night. CD31 is a pan-endothelial adhesion and signaling molecule in the Ig superfamily that is also expressed in platelets and leukocytes^73^. As CD31 is a mechanosensitive protein^74^, alterations in its localization could be linked to circadian changes in blood pressure. Meanwhile, the level of CD31 surface expression affects its propensity to form homophilic versus heterophilic interactions^75^, which could in turn potentiate different forms of interaction between BECs and immune cells.

More broadly, our findings resonate with an emerging recognition of the BBB as a dynamic interface whose functions are adaptively modulated during different physiological states^76^. For example, fasting increases expression of the monocarboxylate transporter MCT1^77^, while shifting mice to a high fat diet transiently decreases brain glucose uptake and GLUT1 levels^78^, suggesting that the BBB responds to metabolic changes. Circadian rhythms have been linked to changes in BBB efflux transport and P-gp transcript levels peak during the mouse active period (night)^55,79^. This has been hypothesized to be a means of saving energy and may also be exploitable for drug delivery strategies^80^. While these and other changes in the BBB are striking, exploration of the underlying mechanisms has largely been limited to changes in total levels of a transcript or protein. The approach described here will allow for investigation into a new dimension of BEC responses to metabolic changes, circadian rhythms, neuronal activity, and other physiological states: changes in the spatial distribution of membrane proteins, which in turn could profoundly affect signaling and transport at the BBB.

### Limitations of the study

We observed that tyrosine-based motifs regulate abluminal localization. However, a gap remains in terms of identifying the downstream trafficking machinery interacting with these motifs; this is an important direction for future work. Based on parallels with epithelial cells, adaptor protein complexes represent promising candidates. Likewise, while we have observed circadian shifts in protein distributions, the mechanisms driving them and the physiological relevance of these shifts remain to be determined.

An additional limitation is that in this approach measurements are restricted areas around the endothelial nucleus, and it has not been formally demonstrated that this region is representative of the full cell surface. Two factors militate for this area being representative of the cell surface overall. First, there is a lack of a diffusion barrier between nuclear and non-nuclear areas of the plasma membrane. In endothelial cells, the tight junction represents a diffusion barrier between the luminal and abluminal plasma membranes, while, likewise, tight junctions create a diffusion barrier between the apical and basolateral domains of epithelial cells. However, there is no known or apparent structural feature that could form such a barrier between nuclear and non-nuclear areas of the endothelial plasma membrane. Second, in some instances, the luminal and abluminal plasma membranes can be partially separated in areas of the cell that do not surround the nucleus (Supplemental Fig.4A,B). This is most apparent in human samples, perhaps owing to their larger dimensions ^81^. In non-nuclear areas of human samples, GLUT1 is observed on the luminal and abluminal membranes, while PODXL is restricted to the luminal membrane, consistent with what is observed in the nuclear area (Supplemental Fig.4C,D). Future developments of this method, potentially incorporating expansion microscopy and automated tracing of the luminal and abluminal membranes, could help to measure a greater area of the endothelial cell surface.

## Methods

### Animals

All procedures conducted were approved by the Harvard Institutional Animal Care and Use Committee (IACUC) and conform to relevant regulatory standards. Mice were maintained on a 12 light/12 dark cycle. Commercially available C57BL/6J (Jackson Laboratory 000664) mice were obtained for this study. A mix of male and female mice were used in this study; no sex-specific effects were noted. Data was collected from adult mice ranging from 8-16 weeks old.

### Human samples

De-identified postmortem brain tissue for this study was obtained at autopsy from Brigham & Women’s Hospital in accordance with the Mass General Brigham Human Research Protection Program (IRB Protocol #2015P001388, D.R.W). The IRB approved a waiver of informed consent for use of de-identified postmortem tissue. Samples of human brain tissue were obtained at autopsy from the mesial temporal lobe cortex of adults, age range 36-72 years old. Exclusion criteria for sample collection were infection at time of death, clinical evidence of neurologic disease, and abnormal brain imaging (CT or MRI brain). Cause of death for all control samples was cardiopulmonary arrest. Post-mortem interval to tissue processing was less than 24 hours for all samples included in this study.

### Cloning and AAV administration

The plasmids used for viral delivery of HA-OAT3, HA-MHC I (wild type and mutants), Alfa-Plasma Membrane and ER-Alfa were produced as follows. Gene fragments were designed based on the murine sequences of OAT3 (UniProt, O88909) or MHC I (Gene, H2-K1, Uniprot P04223). For the Alfa-tag Plasma Membrane construct, plasma membrane labeling was achieved by a fusion protein consisting of an N-terminal SNAP-tag, followed by a GSGSGS linker, then three copies of the Alfa epitope tag^26^, a GSGSGS linker, followed by the CAAX-box membrane targeting sequence from K-Ras isoform 2B (UniProt P01116-2)^25^ at the C-terminus. For the ER-Alfa construct, the ER-targeting sequence of cytochrome p450 2C1 (Uniprot P00180)^25,50^ was placed at the N-terminus, followed by a GSGSGS linker, and four copies of the Alfa tag at the C-terminus. These gene fragments were ordered from Integrated DNA Technologies (IDT) or Twist Bioscience. The sequences of these gene fragments are contained in Supplemental Table 1. Each fragment was Gibson assembled (NEB E2611S) into the pAAV-CAG-SV40NLSf-GFP-3xmiR122-WPRE-HGHpA (Addgene #183775) backbone using the KpnI (NEB R3142S) and EcoRI (NEB R3101S) restriction sites. Custom adeno-associated viruses (AAVs) were produced by the Janelia Viral Tools Team. AAVs were administered intravenously via tail vein injections at a dose of 1x10^12^ viral genomes in adult mice. Brains were harvested for analysis 10 days after administration.

### Immunohistochemistry

Mice were deeply anesthetized with an intraperitoneal injection of ketamine (10 mg/ml)/xylazine (2 mg/ml) in PBS administered at 13 µl/g bodyweight. At 10 minutes, and after confirmation of toe pinch response absence, mice were transcardially perfused with 10 ml of ice-cold 4% paraformaldehyde (PFA), 1% PFA, or PBS (depending on optimal tissue fixation conditions for antibody staining, see Supplemental Table 2) using a peristaltic pump set to a flow rate of 3 ml/min. Brains were post-fixed in 4% PFA, 1% PFA, or methanol for 2 hours or overnight at 4°C with gentle rocking. Samples were then washed with PBS to remove residual PFA or methanol. For cryosections, samples were cryopreserved in a 15% sucrose solution at 4°C overnight until they sank followed by a 30% sucrose solution at 4°C overnight until they sank. Brains were bisected along the midline to obtain sagittal sections. Each hemisphere was frozen in NEG-50 (Richard-Allan Scientific 6502) and sectioned at 30µm on a Leica CM3050 S cryostat. Samples for non-circadian experiments were harvested between 1:30 and 8:30 pm.

Sections were washed in PBS to remove NEG-50, permeabilized in 0.3% PBST (Triton X-100, Sigma-Aldrich T8787-50ML) for 10 mins and then blocked with a 5% normal donkey serum (Jackson Immunoresearch, 017-000-121)/0.3% PBST solution for 1 hour at room temperature. Sections were subsequently incubated with primary antibodies diluted in blocking solution overnight at 4°C, washed 3 times with PBS and incubated with secondary antibodies diluted in blocking solution overnight at 4°C. Primary and secondary antibodies used in this study are listed in Supplemental Table 2.

For the anti-PGP antibody, the permeabilization step was substituted with a 3-minute antigen retrieval step using Trypsin-EDTA at room temperature (Fisher Scientific 25-200-114), PBS wash, brief (< 5 minute) incubation in Dulbecco’s Modified Eagle Media (DMEM; Thermo Fisher 11995065) supplemented with 1% Penicillin/Streptomycin (Thermo Fisher 15140122) and 10% fetal bovine serum (R&D Systems S11150), and a final PBS wash prior to blocking.

For the anti-Cd13 antibody, a different blocking buffer was used - sections were blocked in 1% BSA, 2% Triton-X in PBS overnight at 4°C. Sections were subsequently incubated with primary antibodies diluted in blocking solution for 2 days overnight at 4°C, washed 3 times with PBS and incubated with secondary antibodies diluted in blocking solution for 1 day overnight at 4°C.

Finally, sections were washed 3 times with PBS and once with a 1 mg/ml DAPI stock diluted 1:5000 in PBS (Thermo Fisher Scientific, 62247) for a brief period (4 minutes). For human brain samples, an additional step was included after the above protocol to reduce autofluorescence ^82^: a 0.3% solution of Sudan Black B (Sigma-Aldrich, 199664-25g) in 70% ethanol was added to tissue sections for five minutes, followed by extensive washes with PBS.

Slides were coverslipped (VWR 48393-241) and mounted with ProLong Gold Antifade Mountant (ThermoFisher Scientific, P36934) for imaging.

### Image Acquisition

Representative fluorescent images were acquired using the Abberior GMBH Lightbox software (2024.48.21878-gc86bbd647c) with an Abberior GmBH Facility Line STED microscope on an inverted Olympus IX83 body. This system is equipped with 405 nm, 485 nm, 561 nm, and 640 nm excitation lasers and a pulsed 775 nm STED depletion laser. The 60x/oil immersion lens was used in this study.

#### Selecting laser and filter sets for imaging

For each sample, one membrane marker was visualized using a far-red dye (640 nm excitation line) and the other using a red or orange dye (561 nm excitation line). For images requiring a third channel, a green dye (488 nm excitation line) was used. Prior to imaging, the appropriate dyes (along with DAPI) were chosen on the Lightbox software.

#### STED imaging settings

Images in this study utilized 2D STED. The excitation power used for the STED laser was 2-3X the power used for confocal. STED laser power for markers in mouse tissue was set at 10%. In human tissue, it was set at 3-4%. The specific wavelength power ranged from 1-50% depending on the strength of the antibody and imaging channel (i.e., proteins in the far-red channel required lower power). The MARIX detector, which measures background and STED signal separately thereby minimizing noise, was used for some images. A pixel dwell time of 10-12.5 µs and pixel size of 20-30 nm was used for all images. During imaging and analysis steps, it is important to monitor for signs of smearing of luminal or abluminal membrane signal throughout the nuclear area, as this could indicate z-axis contamination, which could affect measurement reliability.

### Image Quantification

We adapted approaches used to measure polarized protein sorting in epithelial cells and neurons to the endothelial context ^83,84^. Image analysis was performed in ImageJ/Fiji. A freehand line with a width of 25 pixels was used to trace the abluminal and luminal membranes of the BEC. For markers exclusively expressed on one or the other membrane, a co-stain marker was used as a reference for the other membrane. The measure tool was used to determine the mean pixel intensity and length of the traces.

Two methods were utilized to quantify protein distribution: abluminal and luminal percentages and abluminal enrichment scores, which were adapted from previously-described polarity indices.

#### Abluminal and luminal signal percentage

The abluminal/luminal signals were calculated as products of the mean abluminal/luminal intensities and the lengths of the traces along the respective membranes. The total signal was determined as the sum of the abluminal and luminal signal. The abluminal and luminal percentage signals were calculated as follows:

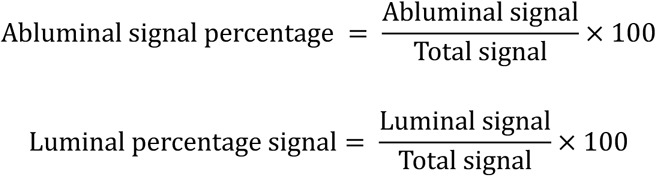

#### Abluminal enrichment score

The abluminal enrichment was calculated as follows:

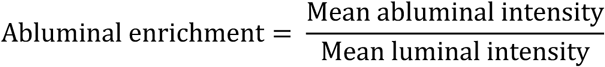

#### Modifications for proteins expressed in pericytes or astrocytes in addition to BECs

For pericyte-expressed proteins, brains were co-stained with integrin β1 or active integrin β1, as well as the pericyte marker CD13 and the BEC marker GLUT1 (Supplemental Fig.1A,B). Areas of the abluminal membrane lacking pericyte coverage were then measured, allowing us to measure the BEC-specific signal (Supplemental Fig.1C). The following steps were followed to demarcate the BEC-only Itgb1 or active Itgb1 signal:

1. A composite with all 3 channels was rendered.
2. A single or multiple traces were drawn on the abluminal membrane where there was overlap with GLUT1 and integrin β1 /Active integrin β1 and no pericyte (Cd13) signal.
3. These traces were saved in the ROI manager.
4. GLUT1 was used as a reference to draw the luminal membrane trace.

To determine if integrin α6 was also localized to astrocyte endfeet in addition to BECs, we co-stained brains with integrin α6 and astrocyte endfoot marker aquaporin-4, or integrin α6 and BEC marker GLUT1 (Supplemental Fig.1D,F). In areas where astrocyte endfeet and BECs were clearly separable, we did not detect overlap between aquaporin-4 and integrin α6 (Supplemental Fig.1D,E). In contrast, we did detect overlap between GLUT1 and integrin α6 (Supplemental Fig.1F,G). These results are consistent with integrin α6 expression in BECs but not astrocyte endfeet.

### Image processing

Images were first post-processed in the Lightbox software. The deconvolve → matrix filter was added to images taken using the MATRIX detector. The deconvolve → intensity filter was added for other images taken using the STED depletion laser. Additional post-processing (e.g., brightness/contrast changes) were done in ImageJ/Fiji^85^.

### Transmission electron microscopy

A wild type mouse was anesthetized as above and perfused with a 5% glutaraldehyde (Electron Microscopy Sciences, 16220), 4% PFA (Electron Microscopy Sciences, 15713-S), 0.4g/mL calcium chloride (Sigma, C7902) solution in 0.1 M sodium cacodylate (Electron Microscopy Sciences, 11653). After dissection, the brain was post-fixed for one hour at room temperature with the same fixative. The brain was then fixed overnight at 4 °C in 4% PFA in 0.1 M sodium cacodylate. After fixation, the brain was washed four times for five minutes per wash in 0.1 M sodium cacodylate. The brain was then cut into 200 µm-thick free-floating sections using a vibratome. Sections were then post-fixed in 1% osmium tetroxide and 1.5% potassium ferrocyanide and dehydrated and embedded in epoxy resin. Ultrathin sections (80 nm) were then cut from the block surface, collected on copper grids, stained with lead citrate and examined under a 1200EX electron microscope (JEOL) equipped with a 2k CCD digital camera (AMT). Electron microscopy services and imaging were performed in the HMS Electron Microscopy Facility.

### Computational search of potential soring motifs in BEC-expressed membrane proteins

We obtained BEC expression data for 15,813 murine genes from an scRNA-seq dataset^14^. Gene names were mapped to protein information in UniProt, including amino acid sequences, lengths, transmembrane and topological domains, and subcellular localization. Focusing on proteins with at least one transmembrane domain and annotated cell membrane localization reduced the set to 1,861 genes. We then screened for YXXØ and [DE]XXXL[LI] motifs in cytoplasmic domains (Ø = V, I, L, F, or M), finding 1,061 genes with YXXØ and 335 [DE]XXXL[LI]. Human orthologs were checked for motif conservations, and genes with moderate-to-high BEC expression yielded 883 YXXØ, 237 [DE]XXXL[LI], and 218 with both motifs. Analyses were performed in Python using numpy, pandas, and regular expression operations. A list of the proteins containing putative motifs are found in Supplemental Table 3. Code for this analysis is available at the following link: https://github.com/gulabneuro/mouse-human-sorting-motifs

### Day versus night measurements

Mice were anesthetized at ZT2 (9am) or ZT14 (9pm), then perfused and processed for immunohistochemistry as described above. Image acquisition, processing and quantification was performed as above by an experimenter blinded to the time when the tissue was harvested. Experimenters were not blinded to experimental groups or protein identity in other experiments.

### Quantification and statistical analysis

Data were analyzed using GraphPad Prism version 9.4.0 software unless otherwise stated, and specific statistical tests are specified in figure legends. Mean abluminal enrichments, percentages, and confidence intervals are listed in Supplemental Table 4. Sample sizes for protein distributions are listed below. PODXL: 5 mice (18, 17, 17,18, and 10 cells measured). Wild type HA-OAT3: 4 mice (13, 10, 13 and 8 cells). Alfa-Plasma Membrane: 3 mice (10, 10 and 11 cells). P-gp: 3 mice (16, 10 and 10 cells). ICAM2: 4 mice (10, 10, 10 and 10 cells). Basigin: 5 mice (9, 10, 10, 10 and 10 cells). CD31: 4 mice (10, 10, 10 and 10 cells). Mfsd2a: 4 mice (11, 10, 10 and 10 cells). GLUT1: 3 mice (11, 10 and 10 cells). TfR: 4 mice (8, 10, 10 and 10 cells). Cav1: 3 mice (10, 10 and 10 cells). Endogenous MHC I: 5 mice (9, 10, 10, 10 and 10 cells). Integrin α6: 3 mice (10, 10 and 10 cells). Integrin β1: 3 mice (10, 10 and 10 cells). Active Integrin β1: 3 mice (10, 10 and 10 cells). HA-OAT3 Y437A/V440E: 4 mice (18, 7, 10 and 8 cells). Wild type HA-H2K1: 4 mice (22, 10, 10 and 10 cells). HA-H2K1 Y342A: 4 mice (15, 10, 10 and 10 cells). GLUT1 human samples: 3 (10, 13 and 10 cells). Cav1 human samples: 3 (10, 12 and 11 cells). PODXL human samples: 3 (10, 10, and 11 cells). Sample sizes for the day/night experiment are as follows. CD31 morning: 6 mice (10 cells in each). CD31 night: 6 mice (10 cells in each). GLUT1 morning: 6 mice (10 cells in each). GLUT1 night: 6 mice (10 cells in each). ICAM2 morning: 6 mice (10 cells in each). ICAM2 night: 6 mice (10 cells in each). PODXL morning: 6 mice (10 cells in each). PODXL night: 6 mice (10 cells in each). Cav1 morning: 6 mice (10 cells in each). Cav1 night: 6 mice (10 cells in each). Statistical analysis in Figure 2 was performed on log-transformed abluminal enrichments; all other tests were performed on raw abluminal enrichments. A p-value <0.05 was considered statistically significant.

As an additional statistical test of the wild type versus tyrosine mutant OAT3 and MHC I comparisons in Figure 3, we generated a linear mixed-effects model using the MATLAB function fitlme with protein (wild type or mutant) as a fixed effect and animal as a random effect. A t-statistic with an absolute value >2 is typically regarded as statistically significant; the values we obtained, as well as p-values <0.05, are consistent with a significant effect of the tyrosine motif mutation on protein distribution and the results of the nested t-tests presented in Figure 3. Model outputs are contained in Supplemental Table 4.

## Acknowledgements

We are grateful to all members of the Gu laboratory for helpful advice and discussion. We appreciate constructive feedback on this manuscript from Jonathan Cohen, Trevor Krolak and Layla Nassar. This work was supported by a T32 Cancer Neuroscience Training Grant 5T32CA272386 (J.A.), the National Institute of Neurological Disorders and Stroke (R35NS116820 to C.G.); C.G. is an investigator of the Howard Hughes Medical Institute.

---

**Supplemental Figure 1.**
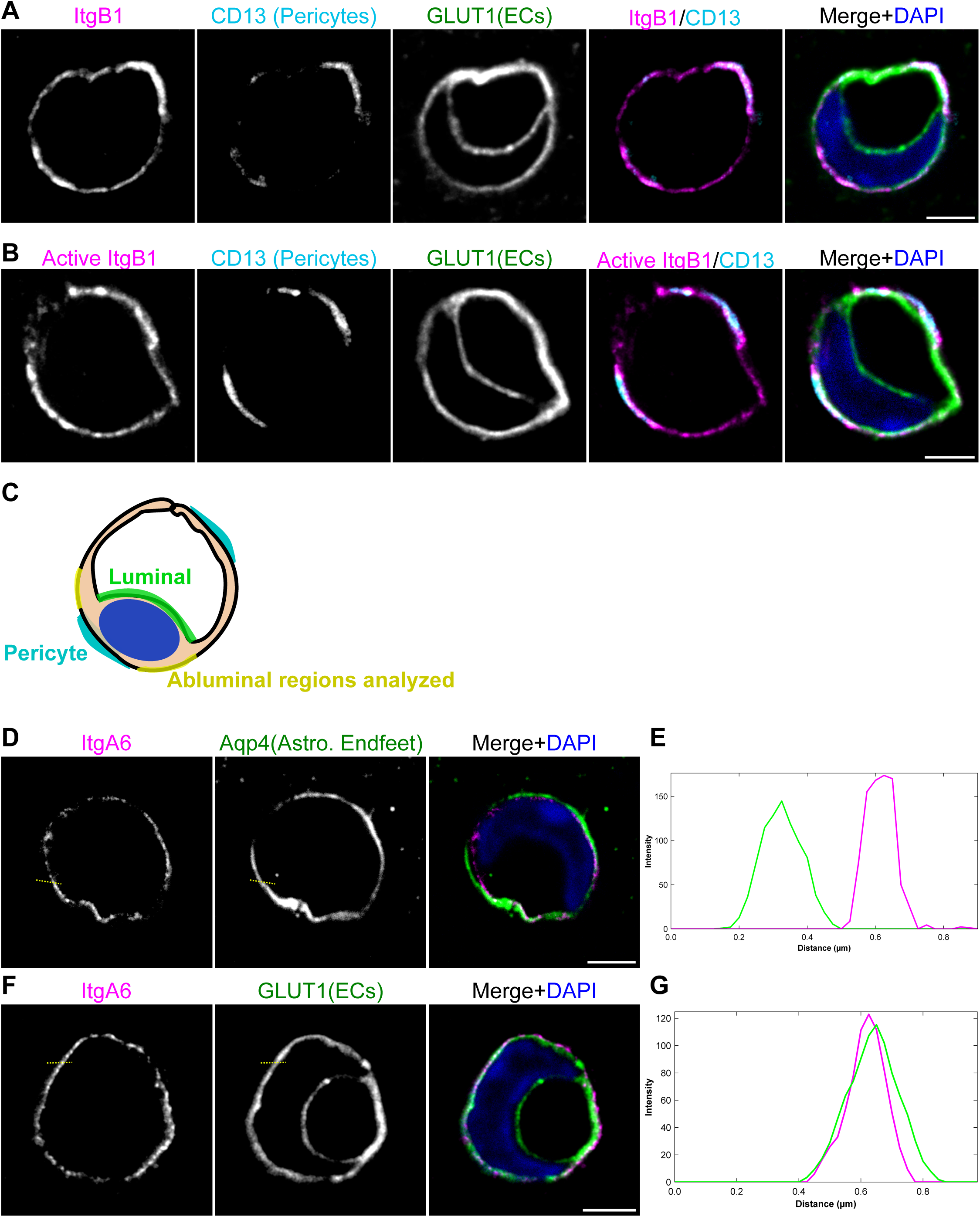
Approach for measuring brain endothelial cell membrane proteins also expressed by pericytes and astrocytes. **(A)** STED images of integrin β1, pericyte-marker CD13 and brain endothelial-cell marker GLUT1. **(B)** STED images of active integrin β1, CD13 and GLUT1. **(C)** Schematic diagram illustrating the approach for measuring abluminal enrichment of proteins expressed by endothelial cells and pericytes. Endothelial luminal (green) and abluminal (purple) plasma membranes, the endothelial nucleus (blue oval) and the position of pericyte processes (cyan) are shown. Example areas of the abluminal plasma membrane without pericyte coverage that would be measured are highlighted in yellow. **(D)** STED images of integrin ⍺6 and astrocyte-endfoot marker Aquaporin-4. **(E)** Intensity profile of (D). Yellow dashed line in (D) indicates the position of the intensity profile. Magenta line represents integrin ⍺6. Green line represents Aquaporin-4. **(F)** STED images of integrin ⍺6 and GLUT1. **(G)** Intensity profile of (F). Yellow dashed line in (F) indicates the position of the intensity profile. Magenta line represents integrin ⍺6. Green line represents GLUT1. Scale bars: 2µm.

**Supplemental Figure 2.**
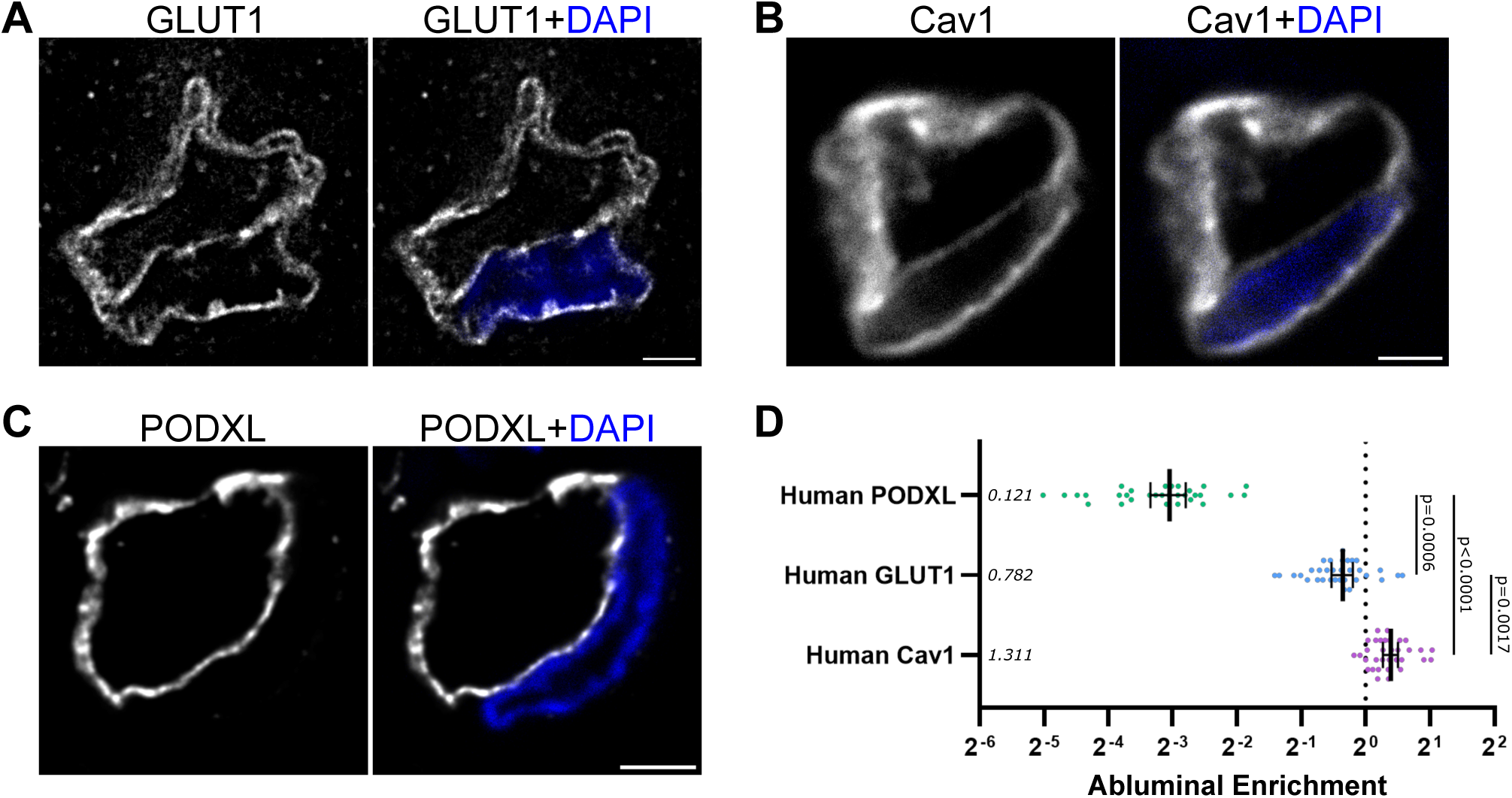
Membrane protein distribution in human brain samples. (A-C) STED images of the indicated proteins in human brain samples with and without DAPI. Scale bars: 2µm. **(D)** Graph of the abluminal enrichments of human Cav1, GLUT1 and PODXL. Data points are measurements from individual cells (mean ± 95% confidence interval). Mean abluminal enrichments are listed. Nested one-way ANOVA with Bonferroni’s multiple comparisons test; p-values indicated.

**Supplemental Figure 3.**
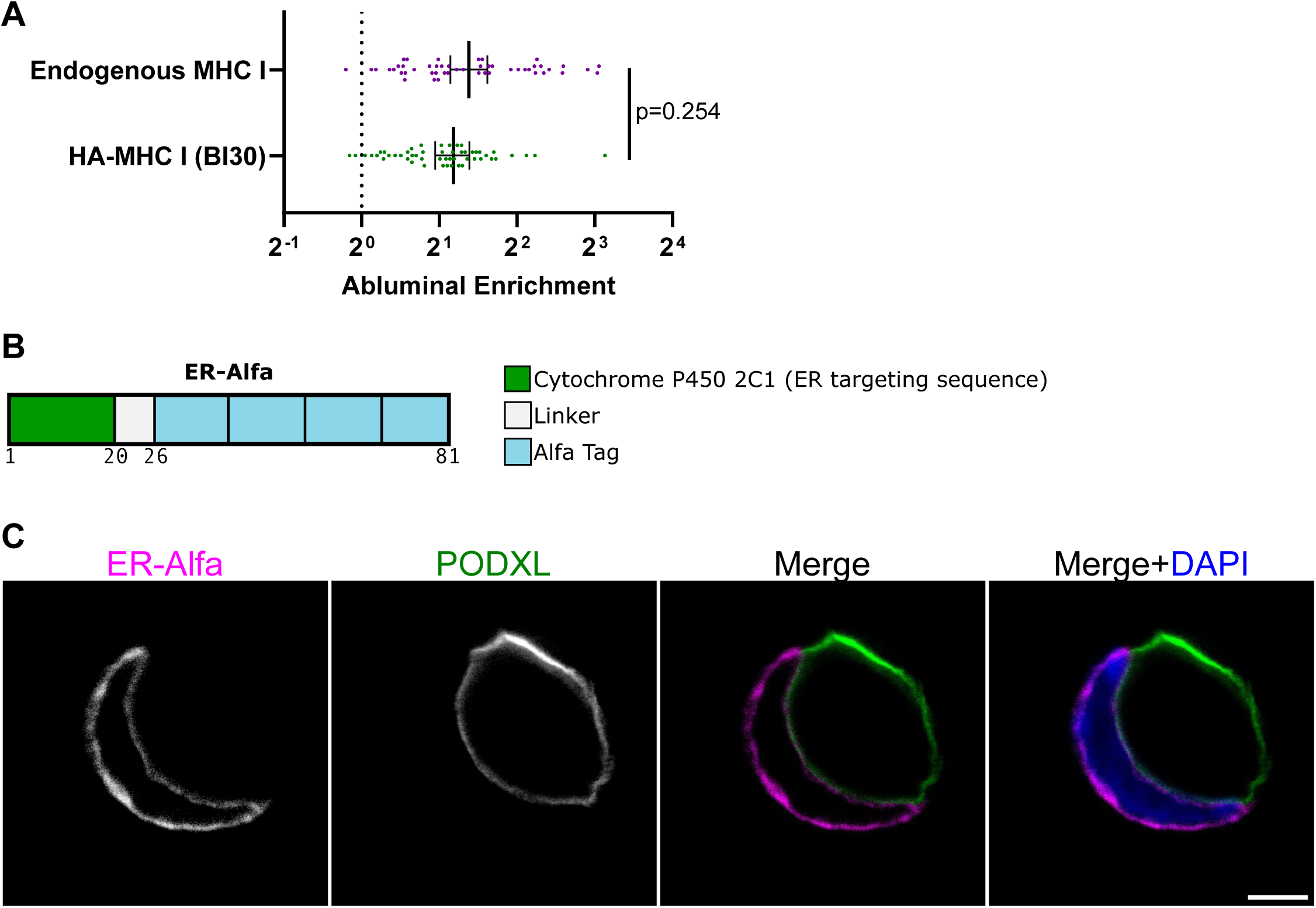
Validation of BI30-expressed constructs. **(A)** Abluminal enrichments of endogenous MHC I and HA-MHC I expressed via BI30 transduction (mean ± 95% confidence interval); p-value from nested t-test indicated. **(B)** Schematic diagram illustrating the construct used to label the endoplasmic reticulum (ER) in BECs. The construct is composed of the ER-targeting sequence from cytochrome p450, a linker, and four copies of the Alfa epitope tag. **(C)** STED image of a brain endothelial cell transduced by AAV-BI30 to express the ER marker from (B). Antibody against the Alfa tag was used to stain for the ER marker; PODXL is an endothelial cell luminal marker. Scale bar: 2µm.

**Supplemental Figure 4.**
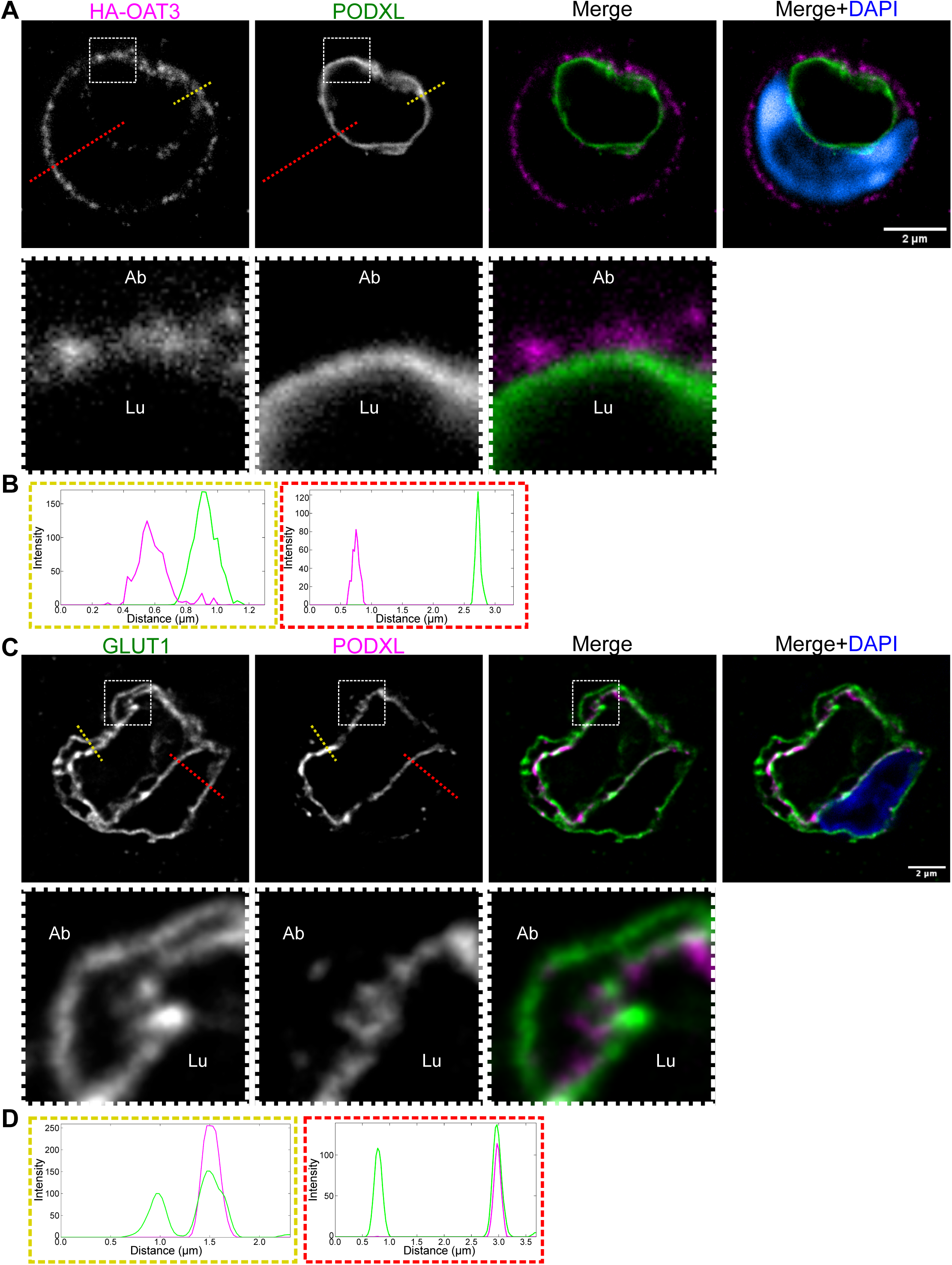
Membrane protein spatial distribution in non-nuclear areas of BECs. **(A)** STED image of an AAV-BI30 transduced endothelial cell in the mouse brain expressing HA-OAT3 co-stained with PODXL. White-dashed box shows inset area. **(B)** Intensity profiles from (A). The yellow (non-nuclear area) and red (nuclear area) dashed lines in (A) indicate the positions of the intensity profiles. Magenta line represents HA-OAT3; green line represents PODXL. **(C)** STED image of GLUT1 and PODXL in human brain tissue samples. **(D)** Intensity profiles from (C). Green line represents GLUT1; magenta line represents PODXL.

## Notes

### Competing Interest Statement

The authors have declared no competing interest.

### Summary of Updates

This updated preprint includes new experiments and data analysis. New experiments and data include an additional protein measured in human data (podocalyxin), an increased animal number in certain experiments, and the results of computational identification of putative sorting motifs. New statistical analyses related to figures 2 and 3 are also included.

## References

1. Amick, J., Gordon, L., and Gu, C. (2026). Cellular and signalling mechanisms that regulate the blood–brain barrier. Nature Reviews Molecular Cell Biology. 10.1038/s41580-026-01000-z.

2. Reese, T.S., and Karnovsky, M.J. (1967). Fine structural localization of a blood-brain barrier to exogenous peroxidase. J Cell Biol 34, 207–217. 10.1083/jcb.34.1.207.

3. Pardridge, W.M. (2007). Blood-brain barrier delivery. Drug Discov Today 12, 54–61. 10.1016/j.drudis.2006.10.013.

4. Lecca, M., Pehlivan, D., Suner, D.H., Weiss, K., Coste, T., Zweier, M., Oktay, Y., Danial-Farran, N., Rosti, V., Bonasoni, M.P., et al. (2023). Bi-allelic variants in the ESAM tight-junction gene cause a neurodevelopmental disorder associated with fetal intracranial hemorrhage. Am J Hum Genet 110, 681–690. 10.1016/j.ajhg.2023.03.005.

5. De Vivo, D.C., Trifiletti, R.R., Jacobson, R.I., Ronen, G.M., Behmand, R.A., and Harik, S.I. (1991). Defective glucose transport across the blood-brain barrier as a cause of persistent hypoglycorrhachia, seizures, and developmental delay. N Engl J Med 325, 703–709. 10.1056/NEJM199109053251006.

6. Guemez-Gamboa, A., Nguyen, L.N., Yang, H., Zaki, M.S., Kara, M., Ben-Omran, T., Akizu, N., Rosti, R.O., Rosti, B., Scott, E., et al. (2015). Inactivating mutations in MFSD2A, required for omega-3 fatty acid transport in brain, cause a lethal microcephaly syndrome. Nat Genet 47, 809–813. 10.1038/ng.3311.

7. Tarlungeanu, D.C., Deliu, E., Dotter, C.P., Kara, M., Janiesch, P.C., Scalise, M., Galluccio, M., Tesulov, M., Morelli, E., Sonmez, F.M., et al. (2016). Impaired Amino Acid Transport at the Blood Brain Barrier Is a Cause of Autism Spectrum Disorder. Cell 167, 1481–1494 e1418. 10.1016/j.cell.2016.11.013.

8. Al-Bachari, S., Naish, J.H., Parker, G.J.M., Emsley, H.C.A., and Parkes, L.M. (2020). Blood-Brain Barrier Leakage Is Increased in Parkinson’s Disease. Front Physiol 11, 593026. 10.3389/fphys.2020.593026.

9. Betz, A.L., and Goldstein, G.W. (1978). Polarity of the blood-brain barrier: neutral amino acid transport into isolated brain capillaries. Science 202, 225–227. 10.1126/science.211586.

10. Worzfeld, T., and Schwaninger, M. (2016). Apicobasal polarity of brain endothelial cells. J Cereb Blood Flow Metab 36, 340–362. 10.1177/0271678X15608644.

11. Sargent, S.M., Bonney, S.K., Li, Y., Stamenkovic, S., Takeno, M.M., Coelho-Santos, V., and Shih, A.Y. (2023). Endothelial structure contributes to heterogeneity in brain capillary diameter. Vasc Biol 5. 10.1530/VB-23-0010.

12. Pfau, S.J., Langen, U.H., Fisher, T.M., Prakash, I., Nagpurwala, F., Lozoya, R.A., Lee, W.A., Wu, Z., and Gu, C. (2024). Characteristics of blood-brain barrier heterogeneity between brain regions revealed by profiling vascular and perivascular cells. Nat Neurosci 27, 1892–1903. 10.1038/s41593-024-01743-y.

13. Kalucka, J., de Rooij, L., Goveia, J., Rohlenova, K., Dumas, S.J., Meta, E., Conchinha, N.V., Taverna, F., Teuwen, L.A., Veys, K., et al. (2020). Single-Cell Transcriptome Atlas of Murine Endothelial Cells. Cell 180, 764–779 e720. 10.1016/j.cell.2020.01.015.

14. Vanlandewijck, M., He, L., Mae, M.A., Andrae, J., Ando, K., Del Gaudio, F., Nahar, K., Lebouvier, T., Lavina, B., Gouveia, L., et al. (2018). A molecular atlas of cell types and zonation in the brain vasculature. Nature 554, 475–480. 10.1038/nature25739.

15. Farquhar, M.G., and Hartmann, J.F. (1957). Neuroglial structure and relationships as revealed by electron microscopy. J Neuropathol Exp Neurol 16, 18–39. 10.1097/00005072-195701000-00003.

16. Korogod, N., Petersen, C.C., and Knott, G.W. (2015). Ultrastructural analysis of adult mouse neocortex comparing aldehyde perfusion with cryo fixation. Elife 4. 10.7554/eLife.05793.

17. Mathiisen, T.M., Lehre, K.P., Danbolt, N.C., and Ottersen, O.P. (2010). The perivascular astroglial sheath provides a complete covering of the brain microvessels: an electron microscopic 3D reconstruction. Glia 58, 1094–1103. 10.1002/glia.20990.

18. Ornelas, S., Berthiaume, A.A., Bonney, S.K., Coelho-Santos, V., Underly, R.G., Kremer, A., Guerin, C.J., Lippens, S., and Shih, A.Y. (2021). Three-dimensional ultrastructure of the brain pericyte-endothelial interface. J Cereb Blood Flow Metab 41, 2185–2200. 10.1177/0271678X211012836.

19. Sabbagh, M.F., and Nathans, J. (2020). A genome-wide view of the de-differentiation of central nervous system endothelial cells in culture. Elife 9. 10.7554/eLife.51276.

20. Kerjaschki, D., Sharkey, D.J., and Farquhar, M.G. (1984). Identification and characterization of podocalyxin--the major sialoprotein of the renal glomerular epithelial cell. J Cell Biol 98, 1591–1596. 10.1083/jcb.98.4.1591.

21. Horvat, R., Hovorka, A., Dekan, G., Poczewski, H., and Kerjaschki, D. (1986). Endothelial cell membranes contain podocalyxin--the major sialoprotein of visceral glomerular epithelial cells. J Cell Biol 102, 484–491. 10.1083/jcb.102.2.484.

22. Kikuchi, R., Kusuhara, H., Sugiyama, D., and Sugiyama, Y. (2003). Contribution of organic anion transporter 3 (Slc22a8) to the elimination of p-aminohippuric acid and benzylpenicillin across the blood-brain barrier. J Pharmacol Exp Ther 306, 51–58. 10.1124/jpet.103.049197.

23. Mori, S., Takanaga, H., Ohtsuki, S., Deguchi, T., Kang, Y.S., Hosoya, K., and Terasaki, T. (2003). Rat organic anion transporter 3 (rOAT3) is responsible for brain-to-blood efflux of homovanillic acid at the abluminal membrane of brain capillary endothelial cells. J Cereb Blood Flow Metab 23, 432–440. 10.1097/01.WCB.0000050062.57184.75.

24. Krolak, T., Chan, K.Y., Kaplan, L., Huang, Q., Wu, J., Zheng, Q., Kozareva, V., Beddow, T., Tobey, I.G., Pacouret, S., et al. (2022). A High-Efficiency AAV for Endothelial Cell Transduction Throughout the Central Nervous System. Nat Cardiovasc Res 1, 389–400. 10.1038/s44161-022-00046-4.

25. Benedetti, L., Marvin, J.S., Falahati, H., Guillen-Samander, A., Looger, L.L., and De Camilli, P. (2020). Optimized Vivid-derived Magnets photodimerizers for subcellular optogenetics in mammalian cells. Elife 9. 10.7554/eLife.63230.

26. Gotzke, H., Kilisch, M., Martinez-Carranza, M., Sograte-Idrissi, S., Rajavel, A., Schlichthaerle, T., Engels, N., Jungmann, R., Stenmark, P., Opazo, F., and Frey, S. (2019). The ALFA-tag is a highly versatile tool for nanobody-based bioscience applications. Nat Commun 10, 4403. 10.1038/s41467-019-12301-7.

27. Schinkel, A.H., Smit, J.J., van Tellingen, O., Beijnen, J.H., Wagenaar, E., van Deemter, L., Mol, C.A., van der Valk, M.A., Robanus-Maandag, E.C., te Riele, H.P., and, et al. (1994). Disruption of the mouse mdr1a P-glycoprotein gene leads to a deficiency in the blood-brain barrier and to increased sensitivity to drugs. Cell 77, 491–502. 10.1016/0092-8674(94)90212-7.

28. de Fougerolles, A.R., Stacker, S.A., Schwarting, R., and Springer, T.A. (1991). Characterization of ICAM-2 and evidence for a third counter-receptor for LFA-1. J Exp Med 174, 253–267. 10.1084/jem.174.1.253.

29. Muramatsu, T. (2016). Basigin (CD147), a multifunctional transmembrane glycoprotein with various binding partners. J Biochem 159, 481–490. 10.1093/jb/mvv127.

30. Lertkiatmongkol, P., Liao, D., Mei, H., Hu, Y., and Newman, P.J. (2016). Endothelial functions of platelet/endothelial cell adhesion molecule-1 (CD31). Curr Opin Hematol 23, 253–259. 10.1097/MOH.0000000000000239.

31. Andreone, B.J., Chow, B.W., Tata, A., Lacoste, B., Ben-Zvi, A., Bullock, K., Deik, A.A., Ginty, D.D., Clish, C.B., and Gu, C. (2017). Blood-Brain Barrier Permeability Is Regulated by Lipid Transport-Dependent Suppression of Caveolae-Mediated Transcytosis. Neuron 94, 581–594 e585. 10.1016/j.neuron.2017.03.043.

32. Nguyen, L.N., Ma, D., Shui, G., Wong, P., Cazenave-Gassiot, A., Zhang, X., Wenk, M.R., Goh, E.L., and Silver, D.L. (2014). Mfsd2a is a transporter for the essential omega-3 fatty acid docosahexaenoic acid. Nature 509, 503–506. 10.1038/nature13241.

33. Dick, A.P., Harik, S.I., Klip, A., and Walker, D.M. (1984). Identification and characterization of the glucose transporter of the blood-brain barrier by cytochalasin B binding and immunological reactivity. Proc Natl Acad Sci U S A 81, 7233–7237. 10.1073/pnas.81.22.7233.

34. Zecca, L., Youdim, M.B., Riederer, P., Connor, J.R., and Crichton, R.R. (2004). Iron, brain ageing and neurodegenerative disorders. Nat Rev Neurosci 5, 863–873. 10.1038/nrn1537.

35. Johnsen, K.B., Burkhart, A., Thomsen, L.B., Andresen, T.L., and Moos, T. (2019). Targeting the transferrin receptor for brain drug delivery. Prog Neurobiol 181, 101665. 10.1016/j.pneurobio.2019.101665.

36. Drab, M., Verkade, P., Elger, M., Kasper, M., Lohn, M., Lauterbach, B., Menne, J., Lindschau, C., Mende, F., Luft, F.C., et al. (2001). Loss of caveolae, vascular dysfunction, and pulmonary defects in caveolin-1 gene-disrupted mice. Science 293, 2449–2452. 10.1126/science.1062688.

37. Razani, B., Engelman, J.A., Wang, X.B., Schubert, W., Zhang, X.L., Marks, C.B., Macaluso, F., Russell, R.G., Li, M., Pestell, R.G., et al. (2001). Caveolin-1 null mice are viable but show evidence of hyperproliferative and vascular abnormalities. J Biol Chem 276, 38121–38138. 10.1074/jbc.M105408200.

38. Lutz, S.E., Smith, J.R., Kim, D.H., Olson, C.V.L., Ellefsen, K., Bates, J.M., Gandhi, S.P., and Agalliu, D. (2017). Caveolin1 Is Required for Th1 Cell Infiltration, but Not Tight Junction Remodeling, at the Blood-Brain Barrier in Autoimmune Neuroinflammation. Cell Rep 21, 2104–2117. 10.1016/j.celrep.2017.10.094.

39. Trevino, T.N., Almousawi, A.A., Martins-Goncalves, R., Ochoa-Raya, A., Robinson, K.F., Abad, G.L., Tai, L.M., Oliveira, S.D., Minshall, R.D., and Lutz, S.E. (2025). A Brain Endothelial Cell Caveolin-1/CXCL10 Axis Promotes T Cell Transcellular Migration Across the Blood-Brain Barrier. ASN Neuro 17, 2472070. 10.1080/17590914.2025.2472070.

40. Aydin, S., Pareja, J., Schallenberg, V.M., Klopstein, A., Gruber, T., Page, N., Bouillet, E., Blanchard, N., Liblau, R., Korbelin, J., et al. (2023). Antigen recognition detains CD8(+) T cells at the blood-brain barrier and contributes to its breakdown. Nat Commun 14, 3106. 10.1038/s41467-023-38703-2.

41. Ayloo, S., Lazo, C.G., Sun, S., Zhang, W., Cui, B., and Gu, C. (2022). Pericyte-to-endothelial cell signaling via vitronectin-integrin regulates blood-CNS barrier. Neuron 110, 1641–1655 e1646. 10.1016/j.neuron.2022.02.017.

42. Yamamoto, H., Ehling, M., Kato, K., Kanai, K., van Lessen, M., Frye, M., Zeuschner, D., Nakayama, M., Vestweber, D., and Adams, R.H. (2015). Integrin beta1 controls VE- cadherin localization and blood vessel stability. Nat Commun 6, 6429. 10.1038/ncomms7429.

43. Shaw, L.M., Messier, J.M., and Mercurio, A.M. (1990). The activation dependent adhesion of macrophages to laminin involves cytoskeletal anchoring and phosphorylation of the alpha 6 beta 1 integrin. J Cell Biol 110, 2167–2174. 10.1083/jcb.110.6.2167.

44. Sonnenberg, A., Linders, C.J., Modderman, P.W., Damsky, C.H., Aumailley, M., and Timpl, R. (1990). Integrin recognition of different cell-binding fragments of laminin (P1, E3, E8) and evidence that alpha 6 beta 1 but not alpha 6 beta 4 functions as a major receptor for fragment E8. J Cell Biol 110, 2145–2155. 10.1083/jcb.110.6.2145.

45. Apodaca, G., Gallo, L.I., and Bryant, D.M. (2012). Role of membrane traffic in the generation of epithelial cell asymmetry. Nat Cell Biol 14, 1235–1243. 10.1038/ncb2635.

46. Guardia, C.M., De Pace, R., Mattera, R., and Bonifacino, J.S. (2018). Neuronal functions of adaptor complexes involved in protein sorting. Curr Opin Neurobiol 51, 103–110. 10.1016/j.conb.2018.02.021.

47. Bonifacino, J.S. (2014). Adaptor proteins involved in polarized sorting. J Cell Biol 204, 7–17. 10.1083/jcb.201310021.

48. Lizee, G., Basha, G., Tiong, J., Julien, J.P., Tian, M., Biron, K.E., and Jefferies, W.A. (2003). Control of dendritic cell cross-presentation by the major histocompatibility complex class I cytoplasmic domain. Nat Immunol 4, 1065–1073. 10.1038/ni989.

49. Kulpa, D.A., Del Cid, N., Peterson, K.A., and Collins, K.L. (2013). Adaptor protein 1 promotes cross-presentation through the same tyrosine signal in major histocompatibility complex class I as that targeted by HIV-1. J Virol 87, 8085–8098. 10.1128/JVI.00701-13.

50. Szczesna-Skorupa, E., and Kemper, B. (2000). Endoplasmic reticulum retention determinants in the transmembrane and linker domains of cytochrome P450 2C1. J Biol Chem 275, 19409–19415. 10.1074/jbc.M002394200.

51. Muller, J.E., Tofler, G.H., and Stone, P.H. (1989). Circadian variation and triggers of onset of acute cardiovascular disease. Circulation 79, 733–743. 10.1161/01.cir.79.4.733.

52. Fabbian, F., Bhatia, S., De Giorgi, A., Maietti, E., Bhatia, S., Shanbhag, A., and Deshmukh, A. (2017). Circadian Periodicity of Ischemic Heart Disease: A Systematic Review of the Literature. Heart Fail Clin 13, 673–680. 10.1016/j.hfc.2017.05.003.

53. Curtis, A.M., Cheng, Y., Kapoor, S., Reilly, D., Price, T.S., and Fitzgerald, G.A. (2007). Circadian variation of blood pressure and the vascular response to asynchronous stress. Proc Natl Acad Sci U S A 104, 3450–3455. 10.1073/pnas.0611680104.

54. Viswambharan, H., Carvas, J.M., Antic, V., Marecic, A., Jud, C., Zaugg, C.E., Ming, X.F., Montani, J.P., Albrecht, U., and Yang, Z. (2007). Mutation of the circadian clock gene Per2 alters vascular endothelial function. Circulation 115, 2188–2195. 10.1161/CIRCULATIONAHA.106.653303.

55. Pulido, R.S., Munji, R.N., Chan, T.C., Quirk, C.R., Weiner, G.A., Weger, B.D., Rossi, M.J., Elmsaouri, S., Malfavon, M., Deng, A., et al. (2020). Neuronal Activity Regulates Blood-Brain Barrier Efflux Transport through Endothelial Circadian Genes. Neuron 108, 937–952 e937. 10.1016/j.neuron.2020.09.002.

56. Zhang, S.L., Yue, Z., Arnold, D.M., Artiushin, G., and Sehgal, A. (2018). A Circadian Clock in the Blood-Brain Barrier Regulates Xenobiotic Efflux. Cell 173, 130–139 e110. 10.1016/j.cell.2018.02.017.

57. Durgan, D.J., Crossland, R.F., and Bryan, R.M., Jr. (2017). The rat cerebral vasculature exhibits time-of-day-dependent oscillations in circadian clock genes and vascular function that are attenuated following obstructive sleep apnea. J Cereb Blood Flow Metab 37, 2806–2819. 10.1177/0271678X16675879.

58. Savolainen, H., Meerlo, P., Elsinga, P.H., Windhorst, A.D., Dierckx, R.A., Colabufo, N.A., van Waarde, A., and Luurtsema, G. (2016). P-glycoprotein Function in the Rodent Brain Displays a Daily Rhythm, a Quantitative In Vivo PET Study. AAPS J 18, 1524–1531. 10.1208/s12248-016-9973-3.

59. Hudson, N., Celkova, L., Hopkins, A., Greene, C., Storti, F., Ozaki, E., Fahey, E., Theodoropoulou, S., Kenna, P.F., Humphries, M.M., et al. (2019). Dysregulated claudin-5 cycling in the inner retina causes retinal pigment epithelial cell atrophy. JCI Insight 4. 10.1172/jci.insight.130273.

60. Lu, T.M., Houghton, S., Magdeldin, T., Duran, J.G.B., Minotti, A.P., Snead, A., Sproul, A., Nguyen, D.T., Xiang, J., Fine, H.A., et al. (2021). Pluripotent stem cell-derived epithelium misidentified as brain microvascular endothelium requires ETS factors to acquire vascular fate. Proc Natl Acad Sci U S A 118. 10.1073/pnas.2016950118.

61. de Caestecker, C., and Macara, I.G. (2024). A size filter at the Golgi regulates apical membrane protein sorting. Nat Cell Biol 26, 1678–1690. 10.1038/s41556-024-01500-0.

62. Shurer, C.R., Levental, K.R., and Levental, I. (2026). Apical and basolateral plasma membranes in epithelial cells have distinct lipidomes and biophysical properties. Proc Natl Acad Sci U S A 123, e2521220123. 10.1073/pnas.2521220123.

63. Armulik, A., Genove, G., Mae, M., Nisancioglu, M.H., Wallgard, E., Niaudet, C., He, L., Norlin, J., Lindblom, P., Strittmatter, K., et al. (2010). Pericytes regulate the blood-brain barrier. Nature 468, 557–561. 10.1038/nature09522.

64. Niewoehner, J., Bohrmann, B., Collin, L., Urich, E., Sade, H., Maier, P., Rueger, P., Stracke, J.O., Lau, W., Tissot, A.C., et al. (2014). Increased brain penetration and potency of a therapeutic antibody using a monovalent molecular shuttle. Neuron 81, 49–60. 10.1016/j.neuron.2013.10.061.

65. Pizzo, M.E., Plowey, E.D., Khoury, N., Kwan, W., Abettan, J., DeVos, S.L., Discenza, C.B., Earr, T., Joy, D., Lye-Barthel, M., et al. (2025). Transferrin receptor-targeted anti-amyloid antibody enhances brain delivery and mitigates ARIA. Science 389, eads3204. 10.1126/science.ads3204.

66. Lobo, A., Launer, L.J., Fratiglioni, L., Andersen, K., Di Carlo, A., Breteler, M.M., Copeland, J.R., Dartigues, J.F., Jagger, C., Martinez-Lage, J., et al. (2000). Prevalence of dementia and major subtypes in Europe: A collaborative study of population-based cohorts. Neurologic Diseases in the Elderly Research Group. Neurology 54, S4–9.

67. Andjelkovic, A.V., Situ, M., Citalan-Madrid, A.F., Stamatovic, S.M., Xiang, J., and Keep, R.F. (2023). Blood-Brain Barrier Dysfunction in Normal Aging and Neurodegeneration: Mechanisms, Impact, and Treatments. Stroke 54, 661–672. 10.1161/STROKEAHA.122.040578.

68. Cirrito, J.R., Deane, R., Fagan, A.M., Spinner, M.L., Parsadanian, M., Finn, M.B., Jiang, H., Prior, J.L., Sagare, A., Bales, K.R., et al. (2005). P-glycoprotein deficiency at the blood-brain barrier increases amyloid-beta deposition in an Alzheimer disease mouse model. J Clin Invest 115, 3285–3290. 10.1172/JCI25247.

69. McCormick, J.W., McCormick, L.A., Chen, G., Vogel, P.D., and Wise, J.G. (2021). Transport of Alzheimer’s associated amyloid-beta catalyzed by P-glycoprotein. PLoS One 16, e0250371. 10.1371/journal.pone.0250371.

70. Engel, J., Jr., Kuhl, D.E., and Phelps, M.E. (1982). Patterns of human local cerebral glucose metabolism during epileptic seizures. Science 218, 64–66. 10.1126/science.6981843.

71. Cornford, E.M., Hyman, S., Cornford, M.E., Landaw, E.M., and Delgado-Escueta, A.V. (1998). Interictal seizure resections show two configurations of endothelial Glut1 glucose transporter in the human blood-brain barrier. J Cereb Blood Flow Metab 18, 26–42. 10.1097/00004647-199801000-00003.

72. Ghosh, C., Westcott, R., Skvasik, D., Khurana, I., Khoury, J., Blumcke, I., El-Osta, A., and Najm, I.M. (2025). GLUT1 and Cerebral Glucose Hypometabolism in Human Focal Cortical Dysplasia Is Associated with Hypermethylation of Key Glucose Regulatory Genes. Mol Neurobiol 62, 10264–10276. 10.1007/s12035-025-04871-z.

73. Newman, P.J., Berndt, M.C., Gorski, J., White, G.C., 2nd, Lyman, S., Paddock, C., and Muller, W.A. (1990). PECAM-1 (CD31) cloning and relation to adhesion molecules of the immunoglobulin gene superfamily. Science 247, 1219–1222. 10.1126/science.1690453.

74. Tzima, E., Irani-Tehrani, M., Kiosses, W.B., Dejana, E., Schultz, D.A., Engelhardt, B., Cao, G., DeLisser, H., and Schwartz, M.A. (2005). A mechanosensory complex that mediates the endothelial cell response to fluid shear stress. Nature 437, 426–431. 10.1038/nature03952.

75. Sun, J., Williams, J., Yan, H.C., Amin, K.M., Albelda, S.M., and DeLisser, H.M. (1996). Platelet endothelial cell adhesion molecule-1 (PECAM-1) homophilic adhesion is mediated by immunoglobulin-like domains 1 and 2 and depends on the cytoplasmic domain and the level of surface expression. J Biol Chem 271, 18561–18570. 10.1074/jbc.271.31.18561.

76. Friedman, A., Prager, O., Serlin, Y., and Kaufer, D. (2025). Dynamic modulation of the blood-brain barrier in the healthy brain. Nat Rev Neurosci. 10.1038/s41583-025-00976-5.

77. Chasseigneaux, S., Cochois-Guegan, V., Lecorgne, L., Lochus, M., Nicolic, S., Blugeon, C., Jourdren, L., Gomez-Zepeda, D., Tenzer, S., Sanquer, S., et al. (2024). Fasting upregulates the monocarboxylate transporter MCT1 at the rat blood-brain barrier through PPAR delta activation. Fluids Barriers CNS 21, 33. 10.1186/s12987-024-00526-8.

78. Jais, A., Solas, M., Backes, H., Chaurasia, B., Kleinridders, A., Theurich, S., Mauer, J., Steculorum, S.M., Hampel, B., Goldau, J., et al. (2016). Myeloid-Cell-Derived VEGF Maintains Brain Glucose Uptake and Limits Cognitive Impairment in Obesity. Cell 165, 882–895. 10.1016/j.cell.2016.03.033.

79. Zhang, S.L., Lahens, N.F., Yue, Z., Arnold, D.M., Pakstis, P.P., Schwarz, J.E., and Sehgal, A. (2021). A circadian clock regulates efflux by the blood-brain barrier in mice and human cells. Nat Commun 12, 617. 10.1038/s41467-020-20795-9.

80. Kim, M., Keep, R.F., and Zhang, S.L. (2024). Circadian Rhythms of the Blood-Brain Barrier and Drug Delivery. Circ Res 134, 727–747. 10.1161/CIRCRESAHA.123.323521.

81. Schaeffer, S., and Iadecola, C. (2021). Revisiting the neurovascular unit. Nat Neurosci 24, 1198–1209. 10.1038/s41593-021-00904-7.

82. Oliveira, V.C., Carrara, R.C., Simoes, D.L., Saggioro, F.P., Carlotti, C.G., Jr., Covas, D.T., and Neder, L. (2010). Sudan Black B treatment reduces autofluorescence and improves resolution of in situ hybridization specific fluorescent signals of brain sections. Histol Histopathol 25, 1017–1024. 10.14670/HH-25.1017.

83. Farias, G.G., Cuitino, L., Guo, X., Ren, X., Jarnik, M., Mattera, R., and Bonifacino, J.S. (2012). Signal-mediated, AP-1/clathrin-dependent sorting of transmembrane receptors to the somatodendritic domain of hippocampal neurons. Neuron 75, 810–823. 10.1016/j.neuron.2012.07.007.

84. Matter, K., Hunziker, W., and Mellman, I. (1992). Basolateral sorting of LDL receptor in MDCK cells: the cytoplasmic domain contains two tyrosine-dependent targeting determinants. Cell 71, 741–753. 10.1016/0092-8674(92)90551-m.

85. Schindelin, J., Arganda-Carreras, I., Frise, E., Kaynig, V., Longair, M., Pietzsch, T., Preibisch, S., Rueden, C., Saalfeld, S., Schmid, B., et al. (2012). Fiji: an open-source platform for biological-image analysis. Nat Methods 9, 676–682. 10.1038/nmeth.2019.

